# Twinkle is not the mitochondrial DNA replicative helicase in *C. elegans*, but may have alternate mitochondrial functions

**DOI:** 10.1101/497305

**Authors:** Hope R. Henderson, Liliya Euro, Anu Suomalainen, Andrew Dillin

## Abstract

Dysfunction of mitochondrial DNA replication machinery is a common cause of mitochondrial diseases. The minimal mammalian replisome is made up of DNA polymerase gamma, replicative helicase Twinkle, and single-stranded DNA binding protein. The replisome is localized to the inner mitochondrial membrane and serves as the site of mitochondrial DNA replication and mitochondrial fission. Recently, a sequence homolog of Twinkle was uncovered in the nematode *Caenorhabditis elegans*. Here, we characterized this homolog, *twnk-1*, and report that *twnk-1* does not function as the primary mitochondrial DNA replicative helicase in this species, as loss of *twnk-1* does not result in reduce mitochondrial DNA levels, or result in other expected mitochondrial dysfunctions such as reduced oxygen consumption rates, increased sensitivity to metabolic perturbations, or reduced muscle function. Instead, *twnk-1* mutants have increased mitochondrial DNA as they age, and exhibit phenotypes associated with mitochondrial stress, including reduced fecundity, an activation of the mitochondrial unfolded protein response, and mitochondrial fragmentation. Our results suggest in *Caenorhabditis elegans*, *twnk-1* does not function as the mitochondrial DNA replicative helicase, but has an alternative function in regulating mitochondrial function.

## INTRODUCTION

Mitochondria are essential organelles, responsible for cellular energy production and the synthesis of metabolites in metazoa. They are the site of β-oxidation of fatty acids and oxidative phosphorylation, have essential roles in calcium homeostasis and apoptosis, and drive anabolic biosynthesis events that support cellular growth. Aberrant mitochondrial function can trigger a wide range of metabolic consequences, and mitochondrial dysfunction is associated with numerous diseases (Gorman *et al.*, 2016).

Mitochondria evolved from archaeabacteria, and became obligate symbionts in many eukaryotic cells. They retain remnants of their evolutionary history; notably, a double membrane and their own circular genome (Friedman and Nunnari, 2014). Over time, most genes from the original bacterial genome have either been transferred to the nuclear genome, or lost. In metazoans, mitochondria maintain a small genome (~15 kb), which contains 12-13 genes crucial for the multimeric electron transport chain (ETC) subunits, while the rest of the ETC proteins are encoded by the nuclear genome. Additionally, the mitochondrial genome contains the sequences to make mitochondrial tRNAs and rRNAs. All other mitochondrial proteins are transcribed from the nuclear genome, and then imported into the organelle by designated protein import mechanisms (Pagliarini *et al.*, 2008; Chacinska *et al.*, 2009).

Of particular interest is the machinery devoted to mitochondrial DNA (mtDNA) replication. Three proteins encoded in the nuclear genome act in concert to make the functional unit of the mtDNA replisome (Fig S1A): (i) DNA polymerase gamma (*POLG* in humans; *polg-1* in *C. elegans*), which elongates mtDNA; (ii) mitochondrial single-stranded binding protein (*mtSSB* in humans; *mtss-1* in *C. elegans*), a protein that binds the lagging strand and stimulates the activity of PolG; and (iii) Twinkle (*TWNK* in humans; *twnk-1* in C. elegans), the mtDNA replicative helicase. Twinkle is a recA/DnaB superfamily helicase, highly similar to the T7 phage replicative helicase, gp4 (Spelbrink *et al.*, 2001; Wanrooij and Falkenberg, 2010). In many metazoan species, including *C. elegans*, Twinkle homologs lack the ancestral DNA binding and nucleotide hydrolysis domains necessary to the primase function. The ancestral N-terminal primase domain is connected by a linker region to the C-terminal helicase domain (Spelbrink *et al.*, 2001; Shutt and Gray, 2006). *In vivo*, Twinkle monomers form a hexamer, making it a ring helicase, as is clearly visible in beautiful electron micrographs from the Solá lab (Fernández-Millán *et al.*, 2015). Human Twinkle unwinds DNA in a 5’ to 3’ direction, hydrolyzing dNTPs at the subunit interface, and leading the replisome complex (Shutt and Gray, 2006). Mutations in PolG and Twinkle cause a variety of disease syndromes, most often affecting the nervous system and muscle (Nunnari and Suomalainen, 2012).

Mutations in PolG and Twinkle cause a variety of documented pathologies. Dominant mutations in PolG and in the Twinkle linker region lead to the accumulation of multiple mtDNA deletions in the muscle, heart, and brain, manifesting as progressive mitochondrial myopathy, sometimes involving sensory neuropathy and parkinsonism (Suomalainen et al. Neurology 1997; Spelbrink et al. Nat Genet 2001). Recessive mutations in PolG and Twinkle, as well as in proteins regulating dNTP pools, results in depletion of mtDNA, often in a tissue-specific manner; recessive mutations in the Twinkle helicase region lead to infantile-onset spinocerebellar ataxia, which leads to severe disability and is fatal within the first two decades of life (Nikali *et al.*, 2005). Our incomplete understanding of the mechanisms of these syndromes, e.g. their tissue-specificity, prevents development of targeted therapies. In fact, there are no treatments for mitochondrial genetic diseases at this time. In order to further understand the molecular basis of these dysfunctions, animal models of genetically-induced mtDNA replication dysfunction are needed.

Over the last decade and a half, there has been an accumulation of evidence suggesting functional conservation of PolG and mtSSB in invertebrate models, including *D. melanogaster* (Thömmes *et al.*, 1995; Williams and Kaguni, 1995; Maier *et al.*, 2001; Baqri *et al.*, 2009; Bratic *et al.*, 2015) and *C. elegans* (Tsang and Lemire, 2002; Bratic *et al.*, 2009; Addo *et al.*, 2010; Bratic, Hench and Trifunovic, 2010). There is also evidence supporting the role of Twinkle as a replicative helicase in *D. melanogaster* (Sanchez-Martinez *et al.*, 2012; Stiban *et al.*, 2014). While a Twinkle homolog has been identified in the *C. elegans* genome (Eki *et al.*, 2007), whether it functions as the mitochondrial DNA replicative helicase is unknown. Here, we sought to determine whether Twinkle is functionally conserved in *C. elegans*.

*C. elegans* possess a mitochondrial genome that is similar in size and structure to the mammalian mitochondrial genome, and shares most of the same genes (Lemire, 2005). Additionally, major metabolic pathways and the major mitochondrial biogenesis pathway are conserved, and similar age-related dysfunctions are observed in mammalian and nematode mitochondria. These include respiratory chain dysfunction, structural abnormalities, decreased oxygen consumption and ATP production, protein aggregation, and accumulation of mtDNA deletions, especially at repetitive sites (Cortopassi and Arnheim, 1990; Tsang and Lemire, 2003; Yasuda *et al.*, 2006; Bratic, Hench and Trifunovic, 2010; Brown, 2012). The conservation of replisome proteins and metabolic aging phenotypes, as well the genetic toolkit and ease of genetic and drug screening in this organism, make *C. elegans* an appealing organism to model mtDNA diseases and to screen for potential drug targets. Thus, we set out to investigate the role of the *C. elegans* Twinkle homolog (Addo *et al.*, 2010).

## MATERIALS AND METHODS

### Strains

The Bristol strain (N2) was used as wild type. The following worm strains used in this study were obtained from the *Caenorhabditis* Genetics Center (University of Minnesota) unless otherwise noted: VC2626 (*F46G11.1(ok3198)X*), VC1224 (*Y57A10A.15(ok1548)/mT1 II; +/mT1* [*dpy-10*(*e128)] II*), MQ887 (*isp-1*(*qm150*)*IV*), TK22 (*mev-1*(*kn1 III*)), CF512 (*rrf-3*(*b26*) *II*; *fem-1*(*hc17*) *IV*), SJ4100 (*zcls13 V*[*hsp-6*::*GFP*]), and SJ410(*zcIs14[myo-3::GFP(mit)]*).

We crossed VC2626 (*F46G11.1(ok3198)X*) with SJ4103 (*zcIs14[myo-3::GFP(mit)]*) to create the homozygozed line used in Figure 6. Backcross of VC2626 (*F46G11.1(ok3198)X*) and cross with SJ4103(*zcIs14[myo-3::GFP(mit)]*) were confirmed with forward primer 5′-CCTTCTCCAAGACTTGACGC-3’ and reverse primer 5′-TACCCAGCGTATTGCACAAG-3’.

### Worm Maintenance and RNAi

Worms were grown on solid agar nematode growth media (NGM) plates at 20°C on OP50 *E. coli* bacteria. For RNAi experiments, worms were fed HT115 *E. coli* bacteria carrying an RNAi construct or an empty construct, L4440, as empty vector control. Synchronized eggs were harvested by timed egg-lay (2 hours) or by bleaching worms with a solution of 1.8% sodium hypochlorite and 0.375 M KOH and arresting L1 animals in M9 buffer (22 mM KH_2_PO_4_ monobasic, 42.3 mM Na_2_HPO_4_, 85.6mM NaCl, 1 mM MgSO_4_) overnight, and grown on RNAi from hatch or L1 arrest. Bacterial feeding in RNAi experiments were conducted from hatch or L1 arrest, as indicated. RNAi strains were taken from the Vidal RNAi library if possible, and from the Ahringer RNAi library otherwise. All clones were sequence-verified.

### Sequence analysis, alignment, and phylogeny

DNA and amino acid sequences were obtained from the National Center for Biotechnology Information (https://www.ncbi.nlm.nih.gov/) and Wormbase (https://wormbase.org/). DNA sequence analysis was performed using National Center for Biotechnology BLAST with standard settings. Amino acid sequence similarity was determined using National Center for Biotechnology BLAST with standard settings. Amino acid sequences were aligned using Clustal Omega software with standard settings, available from the European Bioinformatics Institute (https://www.ebi.ac.uk/Tools/msa/clustalo/). We visualized this alignment using the TCoffee Expresso tool, available at tcoffee.crg.cat. For the phylogenetic tree, sequences were analyzed using Phylodendron, set to create a phenogram with horizontal tree growth with node lengths and inner nodes. This software is available from Indiana University’s Bio-Archive (http://iubio.bio.indiana.edu/treeapp/treeprint-form.html).

### *in silico* modeling of Twinkle

Sequences of Twinkle homologs for alignment and modeling were retrieved from UniProt database. Multiple sequence alignment was done using Promals3D server (http://prodata.swmed.edu/promals3d/). 3D homology models for C-terminal helicase domain of human and *C. elegans* TWINKLE proteins were done in Swiss-Model server using alignment mode. Structural analysis, protein superimposition and figure preparation were done using Discovery Studio v3.5 (BioVia) software.

### Brood Size Assay

Brood size assay performed as described (Chase and Koelle, 2004) with some modifications. Briefly, animals were grown on RNAi from hatch. 10 L4 animals per condition were moved to individual plates, and then moved again every 12 hours. When animals were removed from plates, plates were put at 4° C so eggs would not hatch, and then eggs were counted. Experiment was done in two biological repeats. Statistical significance was calculated using a one-tailed t-test.

### Motility Assay

Motility assay was performed as described (Nawa *et al.*, 2012) with some modifications. Briefly, worms were synchronized by timed egg lay, and grown on RNAi bacteria from hatch. Worms were picked at random from a plate into 50 uL of M9 on an empty plate, and 25 seconds of video was captured immediately. Body bends were counted by eye for each worm; N≥9 worms/strain/experiment were counted. Experiment was done in three biological repeats. Statistical significance was calculated using a two-tailed t-test.

### Oxygen Consumption Rate Measurement

Oxygen consumption rate of whole worms was measured as described (Lin *et al.*, 2016) with some modifications. Briefly, worms were washed from plates and incubated in M9 for 20 minutes to remove residual bacteria. Then, N≥50 worms (10 worms/ well) were transferred to the Seahorse XF96 Cell Culture Microplate (Agilent Technologies, 101085-004) in a total volume of 180 uL. Oxygen consumption rate per well was measured five times using the Seahorse XFe96 Analyzer (Agilent Technologies). Experiment was done in three biological repeats. Statistical significance was calculated using a two-tailed t-test.

### Adult UV stress resistance assays

Resistance of adult nematodes to UVC was performed as described (Chen *et al.*, 2015), with some modifications. After completing egg-laying (D6), animals were treated with 1200 J/m^2^ at D6, and then moved to fresh plates. Irradiation was performed using a CL-1000 Ultraviolent Crosslinker (UVP, CL1000). N≥50 animals per genotype. Survival was scored every 24 hours after. This assay was performed once.

### Larval UV stress resistance assay

Larval UV stress resistance assay was performed as described (Bess *et al.*, 2012), with some modifications. Briefly, animals were bleached, and eggs were left in M9 for 24 hours to achieve L1 arrest. Animals were put on NGM plates without OP50 bacteria, and treated with 10 J/m^2^ at 24, 48, and 72 hours, and then moved to plates with standard OP50 food. Irradiation was performed using a CL-1000 Ultraviolent Crosslinker (UVP, CL1000). Development to L4 or adulthood was scored every 24 hours. N≥33 per condition. Statistical significance was calculated using a one-tailed t-test.

### Whole Animal Microscopy

Synchronized animals were obtained through timed egg lays. For all imaging, unless otherwise noted, animals were D1 adult hermaphrodites and experiments were performed in triplicate with 10–20 animals imaged per experiment. Worms were synchronized by timed egg lay, and grown from hatch on RNAi bacteria when RNAi was used. For fluorescent microscopy, animals from a population were chosen at random under the light microscope. All images were taken using a fixed exposure and gain to just below saturation for the intestinal fluorescence. Images were taken from at least two biological repeats. Animals were anaesthetized with 0.1 M sodium azide (Fisher Scientific, 26628-22-8), and imaged immediately with a Leica MDG41 Stereoscope and Leica DFC3000 G Camera.

### Microscopy of Muscle Mitochondria

Imaging was performed as described (Ichishita *et al.*, 2008), with some modifications. Briefly, synchronized animals were obtained through timed egg lay. For all imaging, animals were D1 adult hermaphrodites and experiments were performed in triplicate with 10–20 animals imaged per experiment. Worms were grown from hatch on RNAi bacteria. Animals were placed directly in M9 media on a glass slide without anesthesia, and imaged immediately (modification courtesy of G. Garcia). Single-plane images of muscle mitochondria from cells just anterior to the vulva were captured in each worm using a Zeiss Oberserver.Z1 Axiovert microscope equipped with a Zeiss axiocam 506 camera, lumencor sola light engine, and ZenBlue software. GFP was visualized by using a 63x/1.4 Plan Aprochromat objective and a standard GFP filter (Zeiss filter set 46 HE). Experiment was performed in more than three biological repeats.

### Electron Microscopy

Samples were prepared as described (Hall, Hartwieg and Nguyen, 2012), with some modifications. Briefly, 200-300 worms were grown on RNAi bacteria from L1 arrest, and then loaded into specimen carriers and fixed using high pressure freezing (Balzers HPM 010 High Pressure Freezer). Samples were then freeze substituted in 1.0% osmium tetroxide, and 0.1% uranyl acetate in acetone at −90°C, and then warmed to −10°C and washed with pure acetone. Worms were embedded in increasing concentrations of Epon resin at room temperature, transferred to flat bottom embedding capsules in pure resin, and cured at 65°C for 48h. Serial sections were cut at 70cnm, and placed onto formvar coated mesh copper grids, and imaged using a FEI Tecnai 12 Transmission electron microscope. Experiment was performed once.

### RNA purification and qRT-PCR

RNA purification were performed as described (Merkwirth *et al.*, 2016), with minor modifications. Briefly, mixed age populations of C*. elegans* were age synchronized by egg bleaching and cultivated on NGM plates. Animals were collected in M9 buffer, centrifuged at 1,000x g for 30 seconds, resuspended in Trizol (Life Technologies, 10296-028), and snap frozen in liquid nitrogen. After several freeze-thaw cycles, total RNA was isolated using chloroform and isopropanol. 1 µg of total RNA was subjected to cDNA synthesis using the QuantiTect Reverse Transcription Kit (Qiagen, 205314). Quantitative real-time PCR reactions were performed with the SYBR Select Master Mix (Life Technologies, 4472920) in Optical 384-well MicroAmp plates (Life Technologies, 4309849) using a QuantStudio 6 Flex (ThermoFisher). *pmp-3* (forward primer 5’-CGGTGTTAAAACTCACTGGAGA-3’, reverse primer 5’-TCGTGAAGTTCCATAACACGA-3’), *cdc-42* (forward primer 5’-AGGAACGTCTTCCTTGTCTCC-3’, reverse primer 5’-GGACATAGAAAGAAAAACACAGTCAC-3’), and Y45F10D.4 (forward primer 5’-AAG CGT CGG AAC AGG AAT C-3’, reverse primer 5’-TTT TTC CGT TAT CGT CGA CTC-3’) were used as housekeeping genes; forward primer 5’-GAT CCT CCA TTG GAT GCT TG-3’ reverse primer 5’-CGG AAG TTT GAT GCC ATT TT-3’ were used to detect *twnk-1*. Experiment was performed once.

### Quantitation and Analysis of Mitochondrial DNA content

Worms (N=10) were lysed in 15 uL of buffer as previously described (Tsang and Lemire, *Biochemica and Biophysica Research*, 2002). The product was diluted 1/1000, and 1 uL of the resulting product was used for each reaction. Absolute quantitation was performed as described (Bratic et al, NAR 2002). Briefly, each experiment included a series of dilutions of pCR2.1 plasmid with the mtDNA NADH dehydrogenase subunit 1 (*nd-1*) sequence inserted. This plasmid was a gift from Dr. A. Trifunovic. Primers for NADH *nd-1* were used to detect the plasmid *nd-1* sequence as well as mtDNA in samples (*nd-1* forward primer is 5′-AGCGTCATTTATTGGGAAGAAGAC-3′ and reverse primer 5′-AAGCTTGTGCTAATCCCATAAATGT-3′). Samples were loaded in triplicate, and absolute quantity of mtDNA was calculated in reference to the Ct values of the plasmid reference. Absolute quantities are displayed as relative to N2 on OP50 or EV RNAi, respectively, at D1. Experiments were done in at least two biological repeats of each experiment. Statistical significance was calculated using a one-tailed t-test.

### Statistical Analysis and Figure Generation

T-tests to analyze significance of data were performed using Sheets (Google). Figures were made using R (RStudio, 1.1.453), GraphPad Prism 6 for Mac OS X (GraphPad Software, Inc.), and Adobe Illustrator CC 22.1 (Adobe).

Plasmids and strains are available upon request. The authors affirm that all data necessary for confirming the conclusions of the article are present within the article and figures, and in supplemental figures and the supplemental table using GSA Figshare.

## RESULTS

### *twnk-1* is the *C. elegans* homolog of Twinkle, and TWNK-1 localizes to mitochondria

Twinkle evolved from the T7 phage gp4 helicase protein, and is found in most eukaryotes, with fungi being a notable exception (Shutt and Gray, 2006). DNA sequence analysis of the *C. elegans* genome confirmed one Twinkle homolog, *F46G11.1*, which we named *twnk-1*. Phylogenetic analysis of Twinkle proteins in other model organisms showed that the nematode branch has significantly diverged from the mammalian branch, as well as from that of other model organisms (Fig 1A). The degree of identity between the human and mouse homologs of Twinkle at the polypeptide level is about 85%; identity between human Twinkle and invertebrate model homologs is about 35-40% (Spelbrink *et al.*, 2001). In *C. elegans* and *C. briggsae*, *twnk-1* carries a 27 amino acids (AA) insertion (Fig 1B).

**Figure 1:**
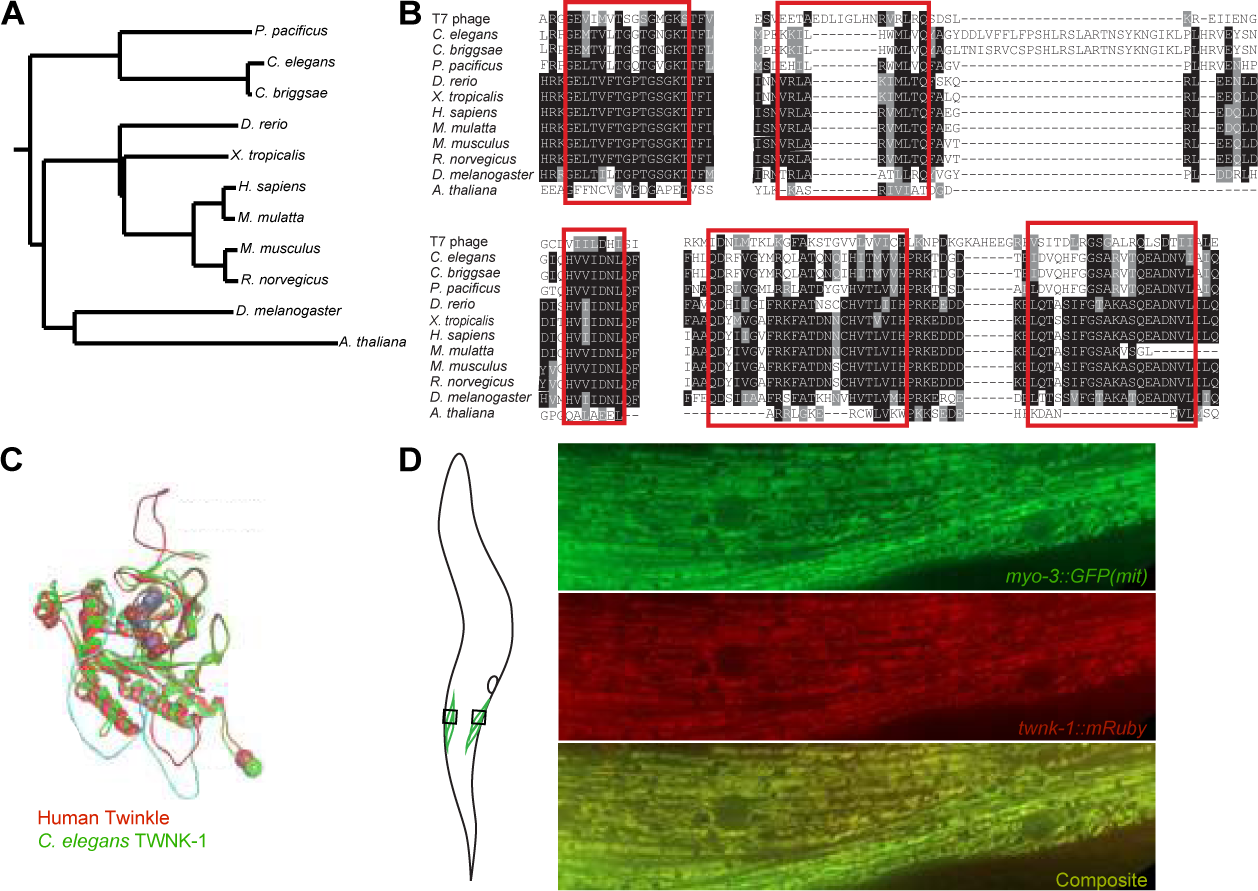
Phylogeny, sequence alignment, and structural analysis of Twinkle homologs. (A) *Phylogenetic tree of twinkle homology:* Phylogenetic tree of model organisms was constructed according to AA sequence of Twinkle homologs or predicted Twinkle homologs. (B) *Sequence alignment of the C-terminal helicase region:* The sequence of the C-terminal helicase region from different species was aligned. Identical (black) and similar (gray) residues in key regions (red boxes) are highlighted. (c) *in silico modeling of Twinkle:* Structure model and overlay of the C-terminal helicase region of human (red) and *C. elegans* (green) Twinkle, with ATP in the Walker A binding cleft. (d) Illustration shows where on the worm body micrographs were taken. Micrographs show localization of TWNK-1::mRuby in myocytes; GFP genetically targeted to muscle mitochondria is used as a mitochondrial marker.

Next, we focused on the functional C-terminal helicase domain. We observed similarities in the amino acid sequence across species in key regions. Overall, the degree of amino acid sequence similarity to the human homolog in *twnk-1* (36% identity) is comparable to that of *polg-1* (37% identity), which is functionally conserved (Tsang and Lemire, 2002; Bratic *et al.*, 2009; Bratic, Hench and Trifunovic, 2010). Our sequence analyses identified no other putative Twinkle homologs or pseudogenes in *C. elegans*. To further examine whether sequence differences would affect the function of the protein, we used *in silico* modeling of the helicase domain to predict its structure. This analysis indicated that the 27 AA insertion is in the first DNA binding loop, likely at the surface where subunits interact (Fig 1C). Despite possible disruptions to the DNA binding, we continued our characterization, as this insertion may have an adaptive role, such as altering the preference to specific DNA binding motifs.

To further confirm *twnk-1* as the functional homolog of Twinkle, we attempted to detect the TWNK-1 protein in mitochondria by quantitative liquid chromatography-mass spectrometry of isolated *C. elegans* mitochondria. We were unable to detect POLG-1 or TWNK-1, suggesting that they exist at levels beneath the detection limits of this method (data not shown). Because of the detection limits, we did not pursue this route of inquiry further. By expressing a TWNK-1 tagged with an mRuby fluorophore under a highly expressed all-tissues promoter (*sur-5p*), we confirmed that the protein localizes to the mitochondria in *C. elegans*, despite lacking a canonical mitochondrial targeting sequence (Fig 1D).

### Loss of *twnk-1* does not deplete mitochondrial DNA

To functionally investigate the role of *twnk-1* in worms, we used RNAi knockdown of *twnk-1*, as well as a knockout strain from the Vancouver consortium (VC2626 (*F46G11.1(ok3198) X*)). In this strain, portions of the N-terminal primase region, the entire linker region, and the Walker A nucleotide binding motif of the C-terminal helicase domain have been deleted (Fig S1B). The resulting truncated mRNA is likely degraded by nonsense-mediated decay (Hug, Longman and Cáceres, 2015). Indeed, we were able to confirm the presence of *twnk-1* transcripts in larval wild type worms, but not *twnk-1* mutant animals (data not shown). We conclude that *twnk-1* is an actively transcribed component of the genome in wild type animals, and that in *twnk-1* mutant animals, no or undetectable levels of *twnk-1* mRNA are allowed to persist.

We predicted that if Twinkle is functionally conserved in *C. elegans*, this strain should suffer from inhibited mtDNA replication. Research in human cell lines confirms that loss of Twinkle is sufficient to deplete mtDNA (Tyynismaa *et al.*, 2004), while Twinkle overexpression in mice is sufficient to increase mtDNA copy number (Tyynismaa *et al.*, 2004; Ylikallio *et al.*, 2010). We examined the effect of loss of *twnk-1* on mtDNA levels in *C. elegans*. Previous work showed that mtDNA content approximately doubles by day 5 of adulthood (D5) in N2 (wild-type) worms. In contrast, when *polg-1* and *mtss-1* were knocked down by RNAi, mtDNA levels on D5 decreased to less than half of their day 1 of adulthood (D1) levels, suggesting functional conservation of these genes with their human homologs (Addo *et al.*, 2010). We replicated these findings, and showing that knockdown of *polg-1* and *mtss-1* significantly decreased the levels of mtDNA by D5, despite similar mtDNA levels on D1. We also tested the effects of knocking down mitochondrial transcription factor A (*TFAM* in humans), a key activator of mitochondrial transcription that is also crucial for mtDNA replication and mtDNA copy number. The worm homolog of TFAM, *hmg-5*, had a similar effect to *polg-1* and *mtss-1*, causing a comparable decrease in mtDNA by D5 (Fig 2A). However, neither RNAi knockdown or knockout of *twnk-1* caused loss of mtDNA in mid-adulthood (Fig 2A, 2B). Furthermore, mtDNA levels of *twnk-1* mutants and animals treated with *twnk-1* RNAi were both higher than the control at D1, and further increased at D5 (Fig 2A, 2B). While these results demonstrate that loss of *twnk-1* does not cause the expected loss of mtDNA as replisome components, *polg-1* and *mtss-1*, the increase in mtDNA upon loss of *twnk-1* may indicate a connection to other mitochondrial functions.

**Figure 2:**
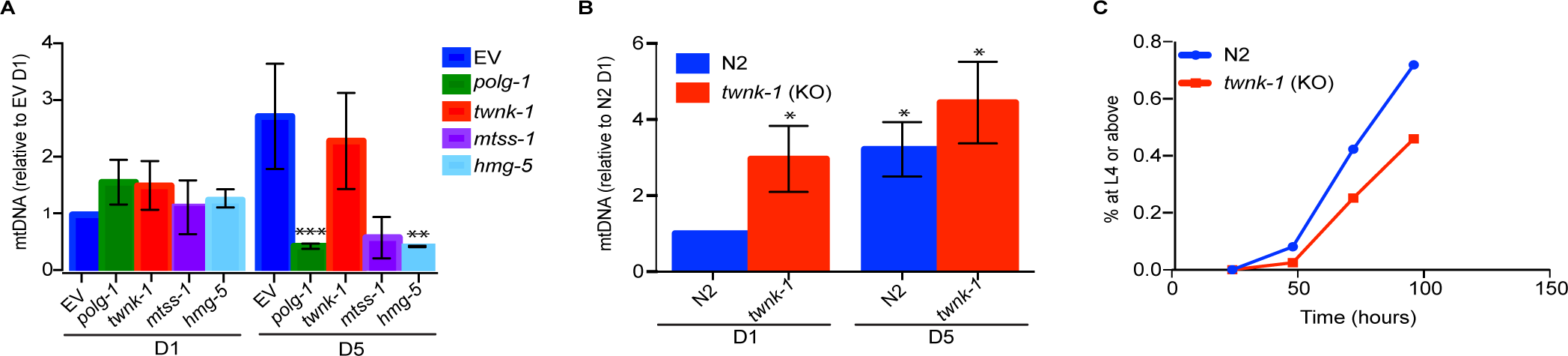
Loss of *twnk-1* increases mtDNA, but decreases fitness under mtDNA replicative stress. (A) *mtDNA levels in twnk-1 knockdown:* Ratio of absolute quantity of mtDNA of animals on replisome component and *hmg-5* RNAi, relative to empty vector at D1, as measured by qPCR. Results shown as mean ± SEM, *** p < 0.001, **p < 0.01, *p < 0.05; ns, p > 0.05. (B) *mtDNA levels in* twnk-1 *mutants:* Ratio of absolute quantity of mtDNA in wild type versus *twnk-1* animals, relative to wild type animals at D1, as measured by qPCR. Results shown as mean ± SEM, *** p < 0.001, **p < 0.01, *p < 0.05; ns, p > 0.05. (C) *Mutation in twnk-1 decreases resistance to UV radiation:* Effect of UV treatment on larval development in wild type and *twnk-1* animals.

To further elucidate the role of *twnk-*1 in mitochondrial DNA replication, we tested the recovery of worms following radiation stress. UV stress in *C. elegans* has been shown to induce mitophagy and loss of mtDNA, and recovery from UV stress requires mtDNA replication (Bess *et al.*, 2012). While we found that adult *twnk-1* animals survived UV stress as well as adult wild type animals (Fig S2), *twnk-1* larval animals were compromised in their ability to recover from UV stress relative to wild type animals, though this effect did not reach statistical significance (Fig 2C). Thus, our results indicate that *twnk-1* function does not dampen normal mtDNA replication or affect UV stress survival in adulthood, but may impair recovery from a stress that requires mtDNA replication during larval development.

### Loss of *twnk-1* does not impair mitochondrial function, but causes stress phenotypes

We next focused on mitochondrial function at the organelle level. If *twnk-1* is functionally conserved, loss of *twnk-1* would be expected to reduce mitochondrial health and metabolic function. To this end, we analyzed oxygen consumption rate (OCR) in early and mid-adulthood. While at individual time points, loss of *polg-1* nor *twnk-1* resulted in significant changes in OCR, the direction of these changes relative to N2 was variable. Taken together, these data suggest that there is no net change in OCR upon loss of *twnk-1* or *polg-1* (Fig 3A and S3A). Conversely, knock down of mitochondrial ribosome protein subunit 5 (*mrps-5*) consistently showed a significant reduction in OCR.

**Figure 3:**
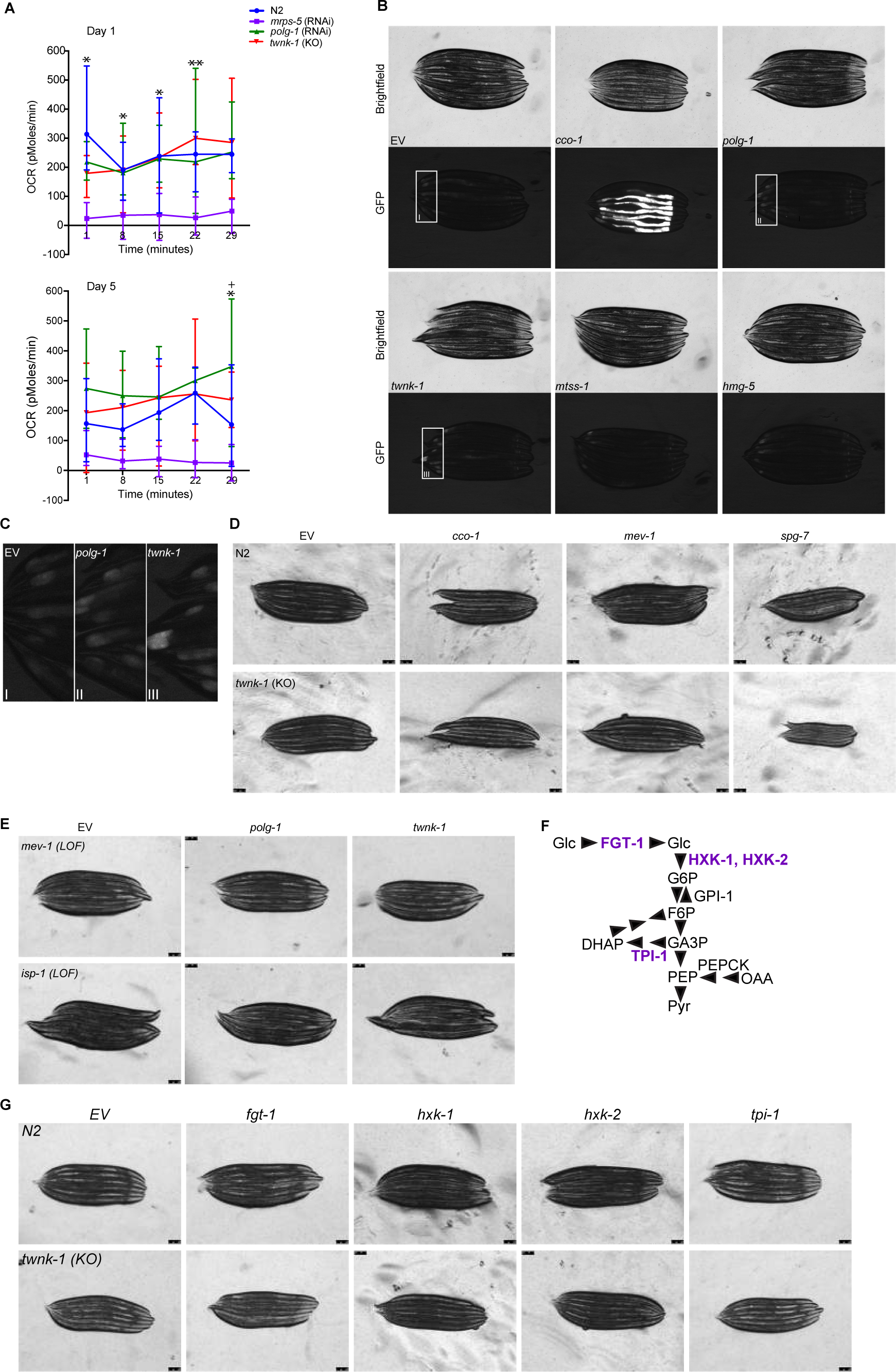
Loss of *twnk-1* causes no gross defects in mitochondrial function. (A) *Mutation of twnk-1 does not affect oxygen consumption rate (OCR):* OCR in *twnk-1* and control *C. elegans* lines was measured using Seahorse XFe96 Analyzer. *mrps-5* was used as a control. Results shown as mean ± SEM. Significance of *twnk-1* relative to age-matched EV animals indicated as follows: *** p < 0.001, **p < 0.01, *p < 0.05; ns, p > 0.05. Significance of *polg-1* (RNAi) relative to age-matched EV animals at the same time point is indicated as follows: +++ p < 0.001, ++p < 0.01, +p < 0.05; ns, p > 0.05. Changes in *mrps-5* (RNAi) OCR were significant for nearly every point. (B) *Knockdown of* twnk-1 *causes a subtle induction of the mitochondrial unfolded protein response (UPR^mt^):* Activation of the UPR^mt^ reporter *hsp-6*::GFP upon RNAi knockdown of replisome components and *hmg-5*. Zoom of hindgut knock down of *polg-1* and *twnk-1*, showing slight upregulation of the *hsp-6::*GFP reporter in (C). (D) *Synthetic interactions with mitochondrial genes*: *twnk-1* and wild type control animals were grown on ETC component (*cco-1*, *mev-1*) and mitochondrial protease (*spg-7*) RNAi. The converse experiment, where ETC mutant animals (*isp-1*, *mev-1*) were grown on replisome RNAi is shown in (E). (F) *Glycolysis flow diagram:* the metabolic pathway from glucose (Glc) to pyruvate (Pyr) is depicted. Glucose transport protein 1 (FGT-1) and the enzymes hexokinase 1 (HXK-1), hexokinase 2 (HXK-2), and triosephosphate isomerase (TPI-1) are highlighted in purple, and knocked-down in wild type and *twnk-1* animals in (G).

Dysfunction at the organelle level may activate the mitochondrial unfolded protein response (UPR^mt^), a protective transcriptional regulation program activated upon mitochondrial stress. In a murine model of Twinkle mutation, critical and conserved UPR^mt^ proteins, including the canonical mitochondrial chaperone protein mtHSP-70 (HSP-6 in *C. elegans*), are upregulated (Khan *et al.*, 2014). Thus, we assayed whether loss of *twnk-1* and other replication-related proteins activated this response by imaging the expression of an *hsp-6*::GFP transcriptional reporter for UPR^mt^ activation. We observed a mild activation of the reporter upon knockdown of *twnk-1*; knockdown of *polg-1*, *mtss-1*, and the TFAM homolog *hmg-5* caused a more subtle induction of this response (Fig 3B, 3C). In case the maternal contribution of these proteins was buffering the animals from dysfunction and stress, we followed animals through two generations of RNAi treatment, but did not see increased reporter activation in the second generation (Fig S3B). Thus, while neither loss of *twnk-1* or disruption to replisome function is a potent activator of the UPR^mt^ in *C. elegans* as assayed by *hsp-6*::GFP, some stress response was evident upon knockdown of *twnk-1*, similar to the level we see caused by expression of expanded repeating glutamates in a *C. elegans* model of Huntington Disease type protein aggregation (Berendzen *et al.*, 2016).

To determine whether *twnk-1* affects metabolic functions of mitochondria, we tested whether loss of *twnk-1* would cause synthetic phenotypes with other perturbations to mitochondrial function. Using RNAi to knockdown electron transport chain (ETC) components, *isp-1* (Complex III) and *cco-1* (Complex IV), we saw no obvious synthetic effects on development, reproduction, or movement in *twnk-1* animals relative to wild-type animals (Fig 3D). In a reciprocal experiment, knockdown of *twnk-1* in ETC complex III mutant, *isp-1*, and ETC complex II mutant, *mev-1*, did not have obvious synthetic effects (Fig 3E). Beyond ETC functions, we hypothesized that oxidative metabolism may be impaired, which would make *twnk-1* mutant animals rely more heavily on glycolysis. To investigate whether *twnk-1* mutants show this metabolic pattern, we knocked down four genes crucial for glycolysis. *twnk-1* mutants showed no gross synthetic effects on development, reproduction, or movement when subject to RNAi knockdown of glucose transmembrane transport gene *fgt-1*, hexokinase genes *hxk-1* and *hxk-2*, or putative triosephosphate isomerase *tpi-1* (Fig 3F, 3G).

Next, we examined mitochondrial morphology, as it may reflect organelle health. Mitochondria in muscles are the most accessible for imaging in worms: in wild-type animals, they exist in lace-like networks through the myocyte. We previously found that overexpressing GFP in mitochondria sensitizes them to fragmentation (unpublished data). Using a strain expressing GFP targeted to mitochondria in muscles, we observed that while mitochondrial networks remained intact in the control, when *polg-1* is knocked down or *twnk-1* is mutated, the mitochondrial network in a subset of muscle cells is fragmented in a stereotypical stress pattern (Fig 4A). The mosaicism of this phenotype is consistent with how muscle dysfunction manifests in human mitochondrial myopathies, where some muscle cells appear normal, and others show loss of ETC function, and disruptions to mitochondrial morphology (Tyynismaa *et al.*, 2005, 2010; Tyynismaa and Suomalainen, 2010; Khan *et al.*, 2014, 2017). To examine the morphology of individual mitochondria in greater detail, we used high magnification transmission electron microscopy on animals at D1 of adulthood (Fig 4B). In diseased patients, mouse models, and *polg-1* mutants, swollen mitochondria with concentric cristae have been observed (Carta *et al.*, 2000; Tyynismaa *et al.*, 2005; Bratic *et al.*, 2009; Nikkanen *et al.*, 2016). We observed no disruptions to mitochondrial cristae upon loss of *polg-1* or *twnk-1*.

**Figure 4:**
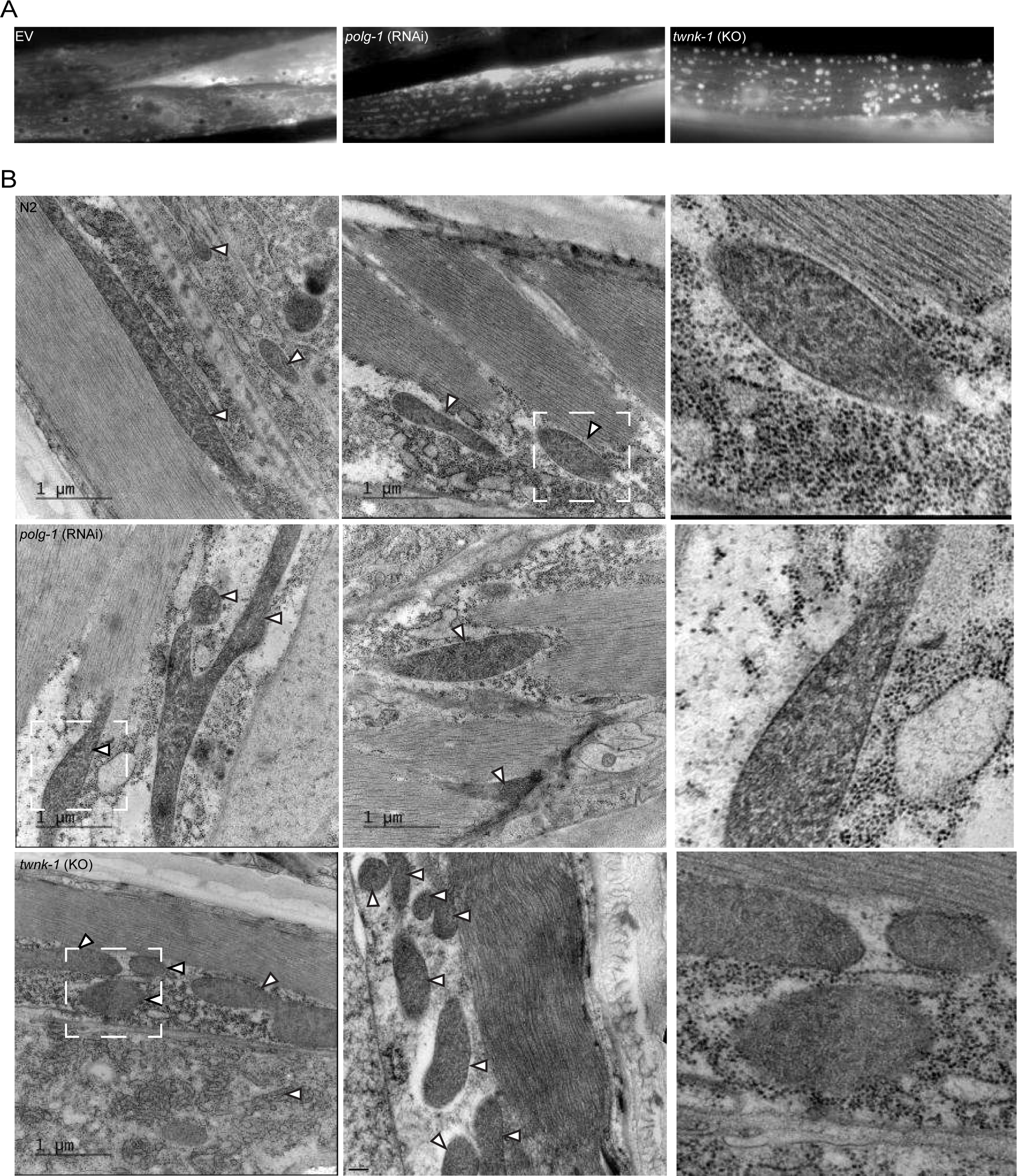
Loss of *twnk-1* causes changes in mitochondrial morphology. (A) *Mutation in twnk-1 causes mitochondrial fragmentation:* Fluorescent imaging of the mitochondrial network in myocytes using GFP targeted to the mitochondria as a mitochondrial marker. Wild-type animals on empty vector (EV) or *polg-1* (RNAi), and *twnk-1* mutant animals were used. (B) *Christae structure appears unperturbed with loss of* twnk-1 *and* polg-1: Electron micrographs of *twnk-1*, *polg-1* (RNAi), and wild-type animals. White arrows point to mitochondria. Images on the right panel are zooms of the corresponding box as indicated by dashed lines.

### Loss of *twnk-1* does not affect development or decrease muscle function, but decreases reproductive capacity

We continued our characterization of loss of *twnk-1* function on organismal health by examining whether *twnk-1* animals exhibited a developmental delay, as mitochondrial biogenesis is crucial for larval development. We found that *twnk-1* animals reached the last larval stage (L4) and the first day of adulthood (D1) concurrent to age-matched wild type worms (Fig 5A). In addition, loss of *twnk-1* showed no synthetic developmental effects with impaired mtDNA replisome function caused by RNAi knockdown of *polg-1* (Fig 5B). The reproductive capacity of the worms was of interest, as *C. elegans* egg production requires robust mitochondrial biogenesis (Tsang and Lemire, 2002). *twnk-1* animals had a reduced brood size relative to N2 animals on empty vector or *polg-1* RNAi (Fig 5C), an effect we also observed informally during strain maintenance. Next, we examined whether loss of *twnk-1* affected muscle function, as loss of mitochondrial function has been linked to decreased muscle function, which worsens with age (Bratic *et al.*, 2009). We found that rate of movement in liquid (thrashing) in early and mid-adulthood was not reduced in *twnk-1* animals (Fig 5D; replicates shown in Fig S5); in fact, these animals maintained their rate of thrashing better than wild type animals in mid-adulthood. This may be related to our finding that *twnk-1* animals have higher mtDNA levels. In summary, loss of *twnk-1* does not seem to negatively impact development and in fact preserves muscle function, but does decrease fecundity.

**Figure 5:**
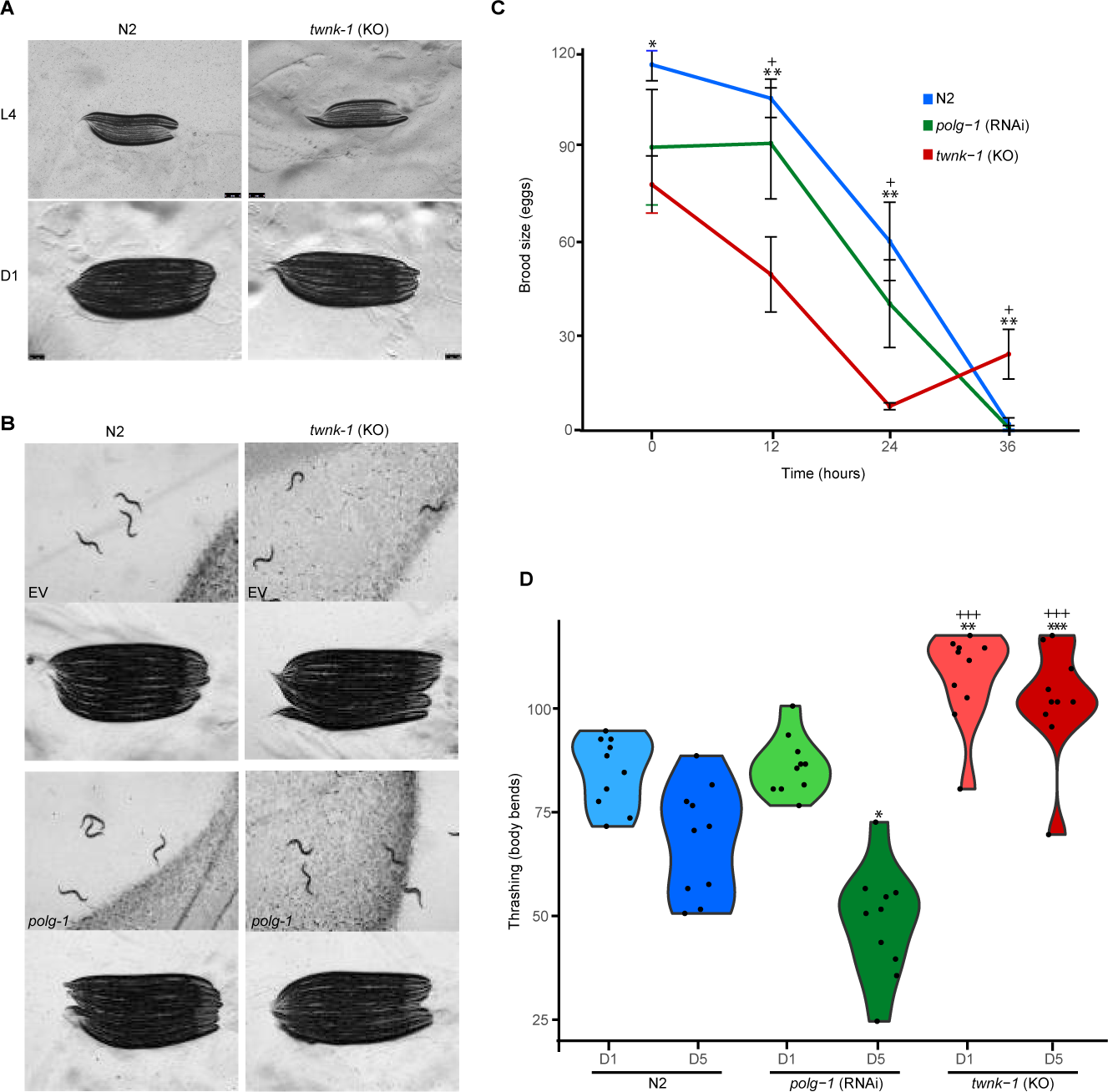
Mutation in *twnk-1* does not impair development, but reduces fecundity. (A) *twnk-1 mutants develop normally: twnk-1* and wild type animals were imaged at age-matched time points in L4 and D1. Note that the development of worms was concurrent. (B) *twnk-1 mutants do not exhibit synthetic effect with* polg-1 *knockdown: twnk-1* and wild type control animals grown on *polg-1* RNAi were imaged. Low magnification pictures show the presence of eggs, indicating that the animals are fertile. (C) *Mutation in twnk-1 reduces fecundity:* Fecundity was assayed by brood size as counted by the number of eggs laid over time for wild type, *polg-1* (RNAi), and *twnk-1* animals. Results shown as mean ± SEM. Significance of *twnk-1* relative to N2 at the same time point is indicated as follows: **p < 0.01, *p < 0.05; ns, p > 0.05. Significance of *twnk-1* relative to *polg-1* at the same time point is indicated as follows: +p < 0.05; ns, p > 0.05. (D) *Animals mutant for twnk-1 increase thrashing:* Motility of wild type, *polg-1* (RNAi), and *twnk-1* animals, as measured by body bends (thrashing) in liquid during early and mid-adulthood. Thrashing measurements were taken over 25 seconds. Significance of *twnk-1* relative to age-matched N2 animals is indicated as follows: *** p < 0.001, **p < 0.01, *p < 0.05; ns, p > 0.05. Significance of *twnk-1* relative to *polg-1* at the same time point is indicated as follows: +++ p < 0.001, ++p < 0.01, +p < 0.05; ns, p > 0.05.

### Screening for the functional homolog of Twinkle

Finally, as our results suggested that *twnk-1* is not functioning as the primary mtDNA replicative helicase in *C. elegans*, we sought to determine if another helicase has this function. We screened a library of RNAi clones, which represents >90% of the known RNA and DNA helicases in *C. elegans* (124 in total). In our primary screen, we looked for synthetic effects on development or fertility in combination with mutations in *twnk-1*, *polg-1*, and ETC component, *isp-1*, with wild type animals and a common sperm-deficient sterile strain as a control. The results of these screens are summarized in Figure 6 (see also Supplementary Table 1). Finally, we screened initial hits and individually selected candidate genes for reduced mtDNA level; this yielded no promising leads for an alternate mtDNA replicative helicase.

**Figure 6:**
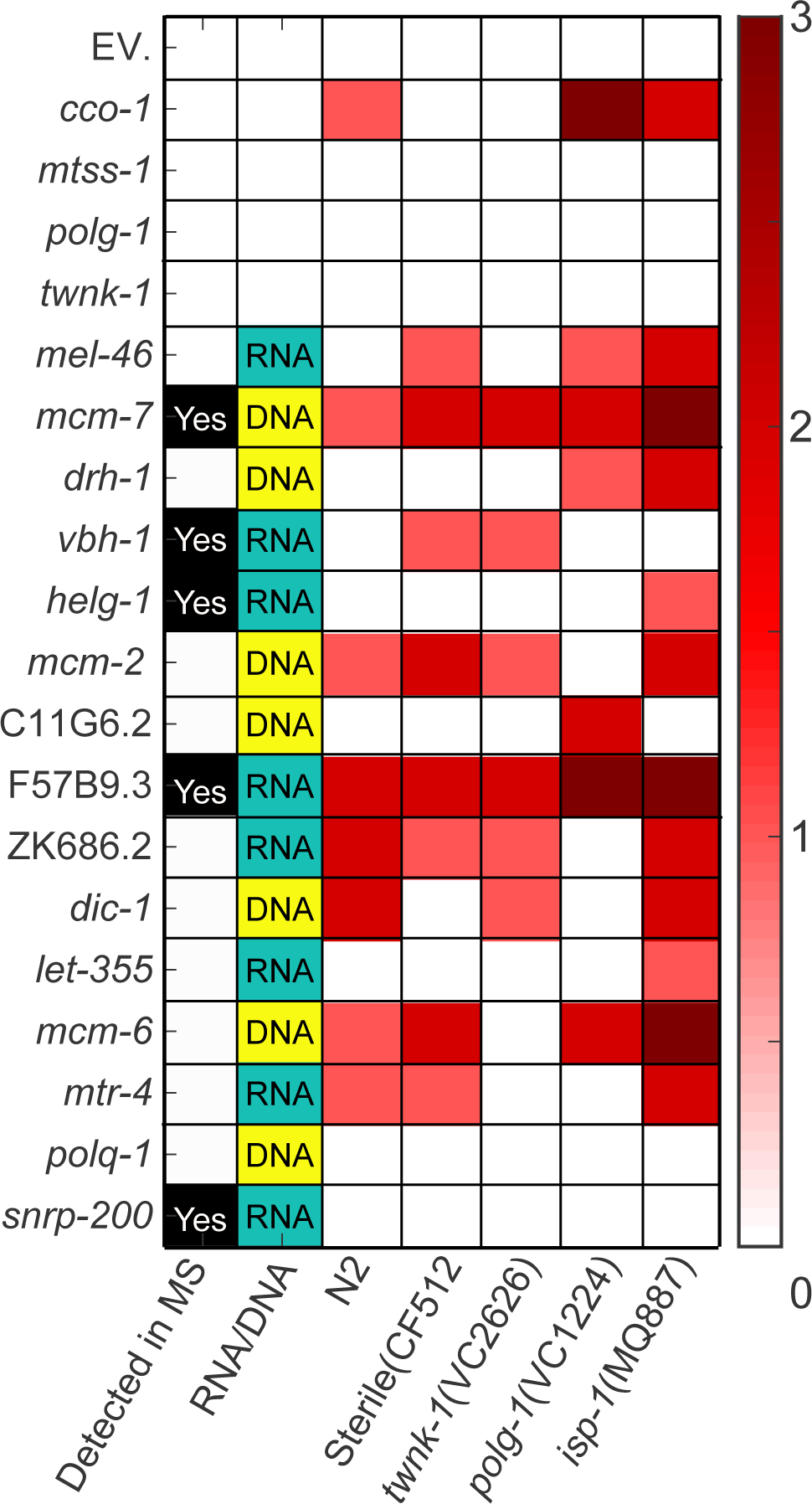
Screening alternative replicative helicases. A summary of the hits from the preliminary screen indicates their features: (i) detection in mass spectrometry of isolated mitochondria (ii) predicted nucleic acid type interacting with the protein, and score (from 0 to 3) for a developmental delay, arrest, and sterility as assessed by microscopy in different genetic backgrounds (iii-vii). See full table of screen results in Table S1.

## DISCUSSION

In this work, we set out to determine whether *twnk-1* is the functional homolog of Twinkle, as an extension of previous works that established conservation of the other integral components of the mitochondrial DNA (mtDNA) replisome. We find, however, that the worm Twinkle homologue *twnk-1* does not function as a replicative helicase for mtDNA, making *C. elegans* an exception in multicellular eukaryotes.

In mammals, PolG, Twinkle, and single-stranded binding protein (mtSSBP) form a conserved minimal mtDNA replisome (Korhonen *et al.*, 2004). In addition to these components, mtDNA maintenance requires the presence of topoisomerases, endonucleases and the transcription factor TFAM (*hmg-5* in *C. elegans*), as well as mitochondrial histones that package the supercoiled mtDNA into mitochondrial chromosomes (Young and Copeland, 2016).

Twinkle and TFAM are the known mtDNA copy number regulators in mammals, the former regulating entry of mtDNA into replication, and the latter binding mtDNA in a histone-like manner, and increasing mtDNA half-life (Tyynismaa et al. Hum Mol Genet 2004; Ylikallio et al. Hum Mol Genet 2010). Knockout of Twinkle and other mtDNA maintenance show that these proteins are essential for embryonic development in mice (Tyynismaa and Suomalainen, 2009). *D. melanogaster* has a conserved Twinkle homolog, which is also an independent regulator of mtDNA copy number (Matsushima and Kaguni, 2007). Twinkle levels also affect replication speed and fidelity in both flies and mice (Ylikallio *et al.*, 2010; Ciesielski *et al.*, 2018). These findings strongly suggest that Twinkle is functioning as a conserved mtDNA replicative helicase, licensing mtDNA to enter replication in both vertebrate and invertebrate metazoans. In the yeast *S. cerevisiae*, however, there is no Twinkle homologue, indicating its later evolutionary emergence, or loss in this organism (Sedman *et al.*, 2005).

Analyzing the TWNK-1 amino acid sequence in *C. elegans* revealed a similar degree of conservation to the human homologue as was observed between the human and worm PolG, which is functionally conserved. We were further encouraged by the *in silico* modeling of the structure of TWNK-1, which indicated similarity in the C-terminal helicase domain, and the localization of TWNK-1::mRuby to the mitochondria. However, animals with knockdown or knockout of *twnk-1* were viable and did not show loss of mtDNA or decreased oxygen consumption, contrary to the detrimental phenotypes of knock-down of other replisome components. These results indicate that *twnk-1* lacks functional conservation with the human homolog.

In fact, upon loss of *twnk-1*, mtDNA was slightly, but significantly, elevated, suggesting that *twnk-1* may functions in non-replicative mtDNA processing, such as repair, or that increased mtDNA is the result of a secondary response, such as mitochondrial biogenesis spurred by stress. Following UV stress in *C. elegans*, mitochondria and mtDNA are degraded by mitophagy. In order to recover from this acute stress, cells undergo a burst of mitochondrial biogenesis and mtDNA replication (Bess *et al.*, 2012). *twnk-1* mutant adults were not sensitive to UV stress, but worms in the first larval stage (L1) had reduced resistance to UV stress. These results suggest that *twnk-1* may have acquired a role as a repair-or stress-specific helicase during development. *twnk-1* mutants also show reduced fecundity. As oocyte production requires a surge in mitochondrial biogenesis (Tsang and Lemire, 2002), this result may be associated with a deficiency in some kind of mitochondrial function.

The hypothesized genetic interaction of *twnk-1* with different cellular pathways, including other replisome components (*polg-1*), glycolysis (*hxk-1*, *hxk-2*, *fgt-1*, *tpi-1*), and oxidative phosphorylation (*isp-1*, *mev-1*, *cco-1*), revealed no synthetic developmental or physiological defects, which is consistent with *twnk-1* not having a role in respiratory chain function or nutrient metabolism. Interestingly, knocking down *spg-7*, the *C. elegans* homolog of the mitochondrial quality control protease paraplegin, caused developmental arrest of the *twnk-1* mutant. *spg-7* is the *C. elegans* homolog of the mitochondrial quality control protease, parapalegin; its synthetic interaction with *twnk-1* may suggest a prior challenge to the mitochondrial protein landscape, or indicate that *twnk-1* is related to managing mitochondrial stress. Finally, imaging of muscle mitochondria revealed that knockdown of *twnk-1* causes the mitochondrial network to fragment, a classic indicator of mitochondrial stress in *C. elegans* (Ichishita *et al.*, 2008). Taken together, our results indicate that TWNK-1 has a pleiotropic role in regulating mitochondrial form and function distinct from mtDNA replication in *C. elegans*.

An indication for possible such role may be a finding in human cell lines, where Twinkle maintains its punctate mitochondrial localization and association with the inner mitochondrial membrane (IMM) even in cells lacking mitochondrial DNA (P_0_); this is in marked contrast to other replisome proteins, which are diffuse in the absence of mtDNA (Spelbrink *et al.*, 2001; Rajala *et al.*, 2014; Gerhold *et al.*, 2015). Specifically, human Twinkle was found to associate with cholesterol in the IMM (Gerhold *et al.*, 2015). Additionally, mammalian cell culture models show that constriction of the IMM is an important early step in mitochondrial fission (Cho *et al.*, 2017), and preferentially occurs where the replicating mtDNA nucleoid is located (Lewis, Uchiyama and Nunnari, 2016). The morphological and physiological defects we observed upon loss of *twnk-1* may imply that, in *C. elegans*, Twinkle serves as an IMM protein that is important to mitochondrial membrane structure, and organelle morphology and dynamics. While research on Twinkleopathies has focused on loss of replicative functions, our findings suggest that it may be worth investigating whether other kinds of mitochondrial distress are also causative in these syndromes. We hoped to examine the effect of expressing human Twinkle in *C. elegans*, but were unable to confirm successful overexpression of human Twinkle in *C. elegans*, or generate viable animals expressing a dominant mutation in the linker region (PEO type) of human Twinkle (data not shown).

We conclude that *C. elegans* is an evolutionary divergence from the typical metazoan mtDNA replisome, having recruited the conserved Twinkle protein to alternative mitochondrial tasks. Finally, we screened dozens of other helicases but were unable to find one that appeared to be functioning as an alternative primary mtDNA replicative helicase. Investigations of additional genes may find a functional homolog of Twinkle in *C. elegans*, and set the stage for modeling mtDNA diseases in this organism.

## AUTHOR CONTRIBUTIONS

A.S., A.D., and H.R.H. conceived of the project. A.S. and A.D. provided ongoing intellectual input and support for the project. J.J.M. did mass spectrometry experiments and analyzed mass spectrometry data. L.E. did *in silico* modeling of human and *C. elegans* Twinkle homologs; H.R.H did all other experimental work. H.R.H. wrote the manuscript, with help from A.S..

## ACKNOWLEDGEMENTS

We thank Dr. S. C. Wolff for intellectual guidance during early stages of the project, and assistance with the helicase screen,. We thank Dr. J. Durieux for creation of the mRuby::Twinkle worm strains, and the illustration in Fig 1D. We thank Dr. J. J. Moresco for mass spectrometry. We thank M. Sanchez for electron microscopy. We thank A. Zhelokhovtseva for help with the helicase screen. We thank Dr. R. Bar-Ziv for help with figure and manuscript preparation.

Howard Hughes Medical Institute and the National Institutes of Health (R37AG024365) supported this work. H.R.H. was supported by a National Science Foundation Graduate Research Fellowship. Some of the nematode strains used in this work were provided by the *Caenorhabditis* Genetics Center (University of Minnesota), which is supported by the NIH– Office of Research Infrastructure Programs (P40 OD010440). This work used the Vincent J. Coates Genomics Sequencing Laboratory at UC Berkeley, supported by NIH S10 Instrumentation Grants (S10RR029668 and S10RR027303).

A.D. is a cofounder of Proteostasis Therapeutics, Inc. and Mitobridge, Inc. and declares no financial interest related to this work.

## SUPPLEMENTARY FIGURE LEGENDS

**Supplementary Figure 1:**
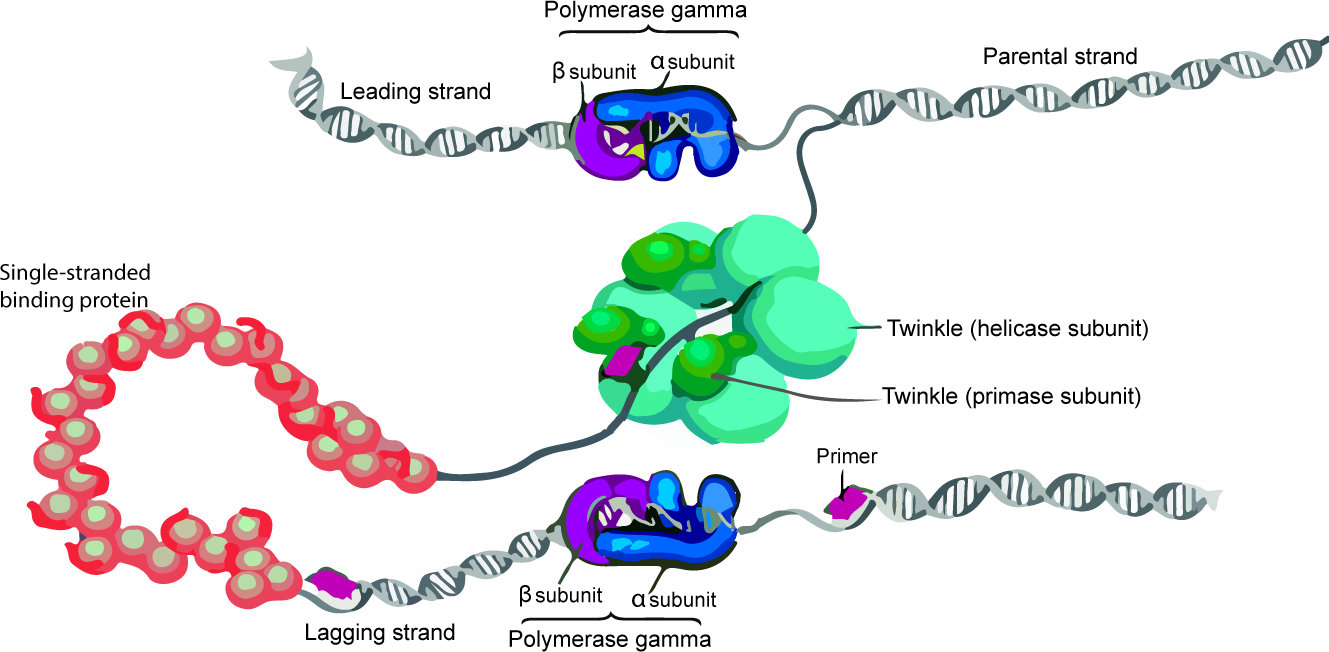
Schematics of mtDNA replisome and *twnk-1* locus mutation. (A) Schematic of the minimal mitochondrial DNA replisome in characterized metazoans. (B) Schematic of *twnk-1* locus in wild type versus *twnk-1* mutant.

**Supplementary Figure 2:**
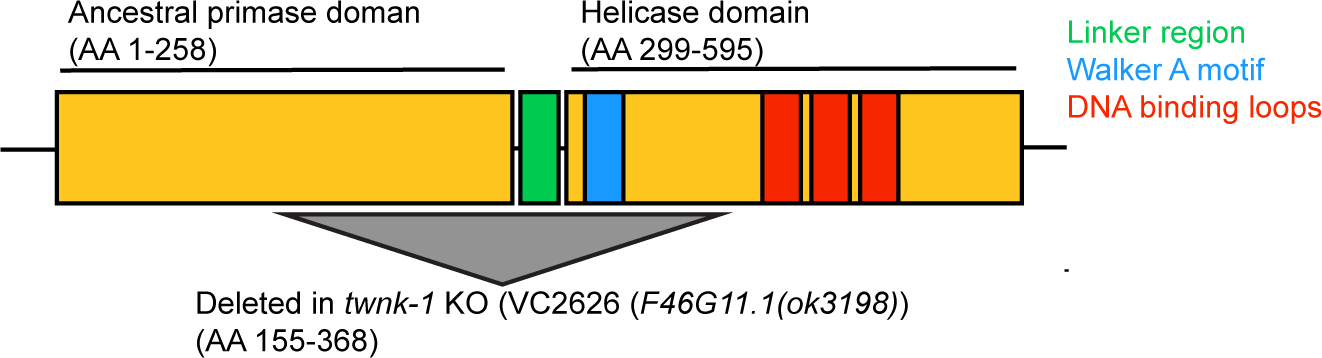
*twnk-1* adults are not sensitized to UV stress. Survival curve of adult wild type versus *twnk-1* mutants after UV treatment.

**Supplementary Figure 3:**
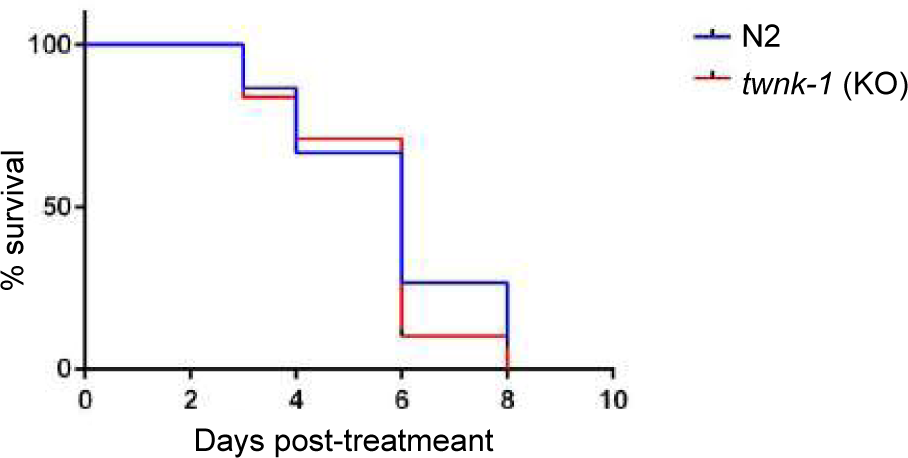
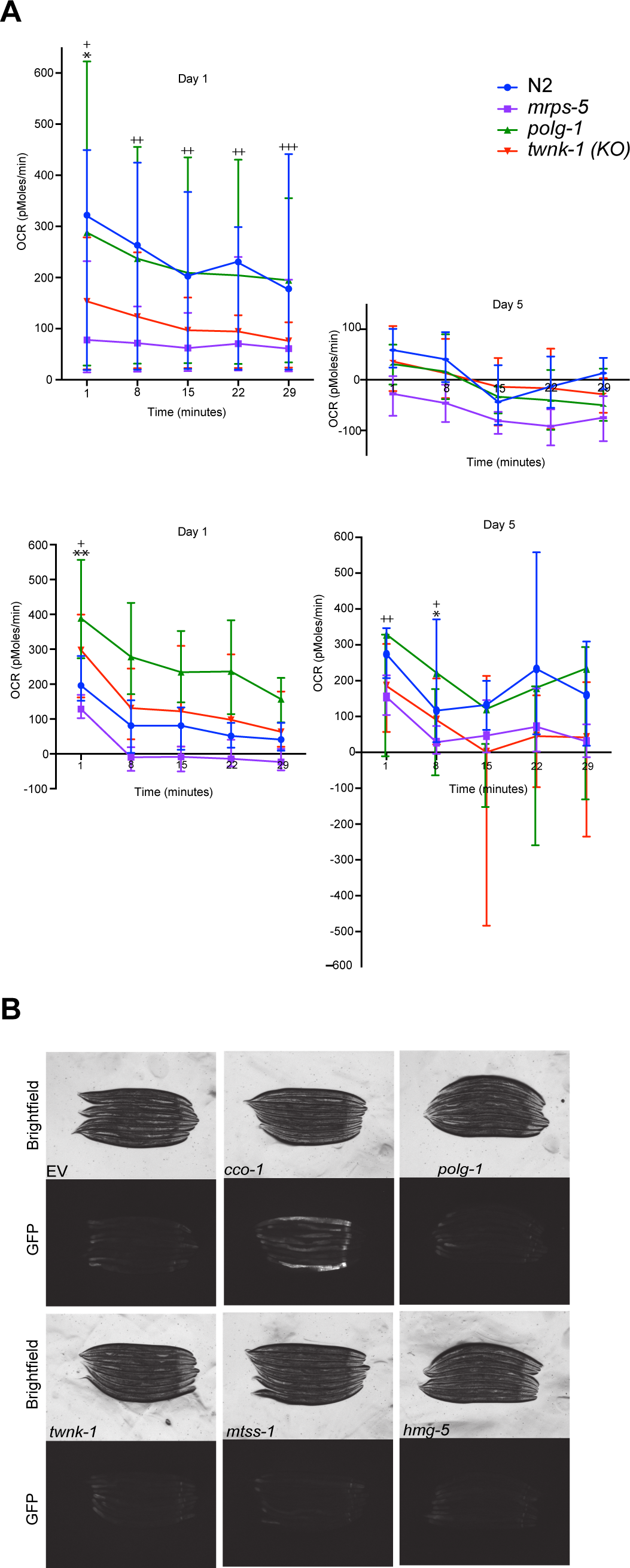
*twnk-1* does not decrease OCR, but does cause subtle activation of the UPR^mt^. (A) *Mutation of* twnk-1 *does not affect oxygen consumption rate (OCR):* OCR in *twnk-1*, *polg-1* (RNAi), and control *C. elegans* lines was measured using Seahorse XFe96 Analyzer. Two independent replicates are shown here; results shown as mean ± SEM. Significance of twnk-1 relative to wild type indicated by: *** p < 0.001, **p < 0.01, *p < 0.05; ns, p > 0.05. Significance of *polg-1* relative to wild type indicated by: +++ p < 0.001, ++p < 0.01, +p < 0.05; ns, p > 0.05. B) *Two generations of RNAi knock down of twnk-1 causes a subtle induction of the mitochondrial unfolded protein response (UPR^mt^):* Activation of the UPR^mt^ reporter *hsp-6*::GFP upon two generations of RNAi knockdown of replisome components and *hmg-5*.

**Supplementary Figure 4:**
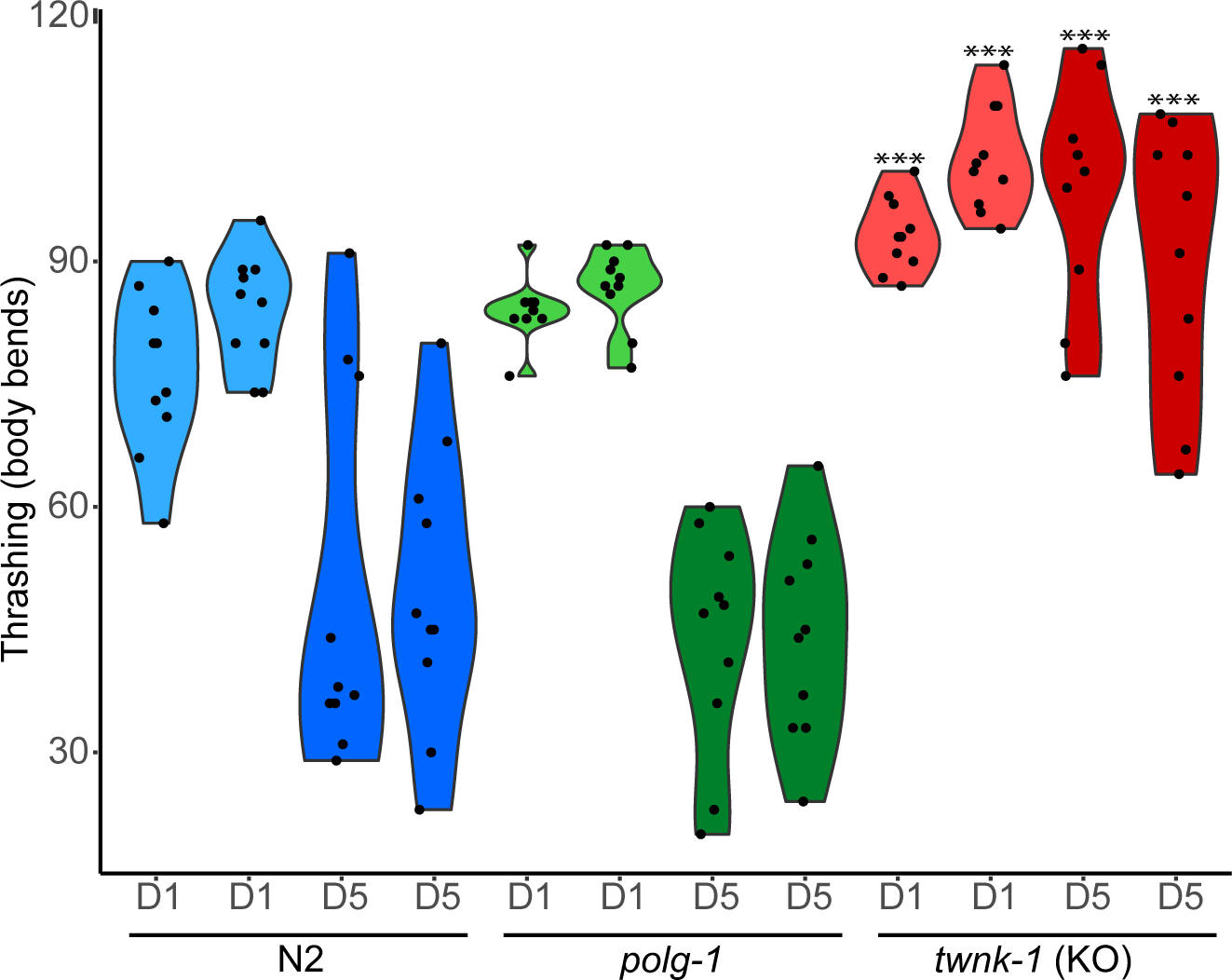
Animals mutant for *twnk-1* show increased thrashing. Motility of wild type, *polg-1* (RNAi), and *twnk-1* animals, as measured by body bends (thrashing) in liquid during early and mid-adulthood. Two independent replicates are shown here. Results shown as mean ± SEM; significance relative to age-matched N2 animals shown as *** p < 0.001, **p < 0.01, *p < 0.05; ns, p > 0.05.

**Supplementary Table 1:**
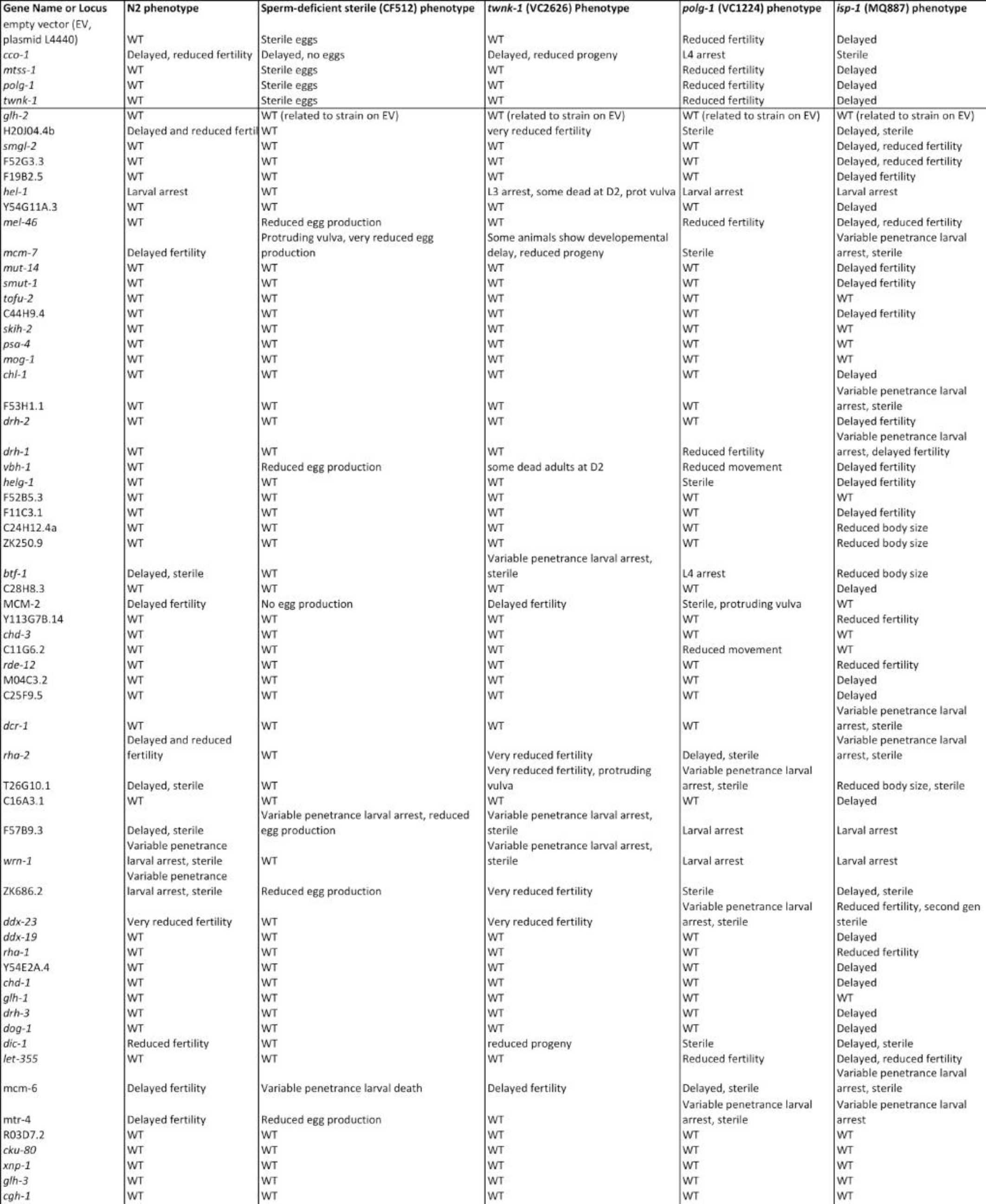
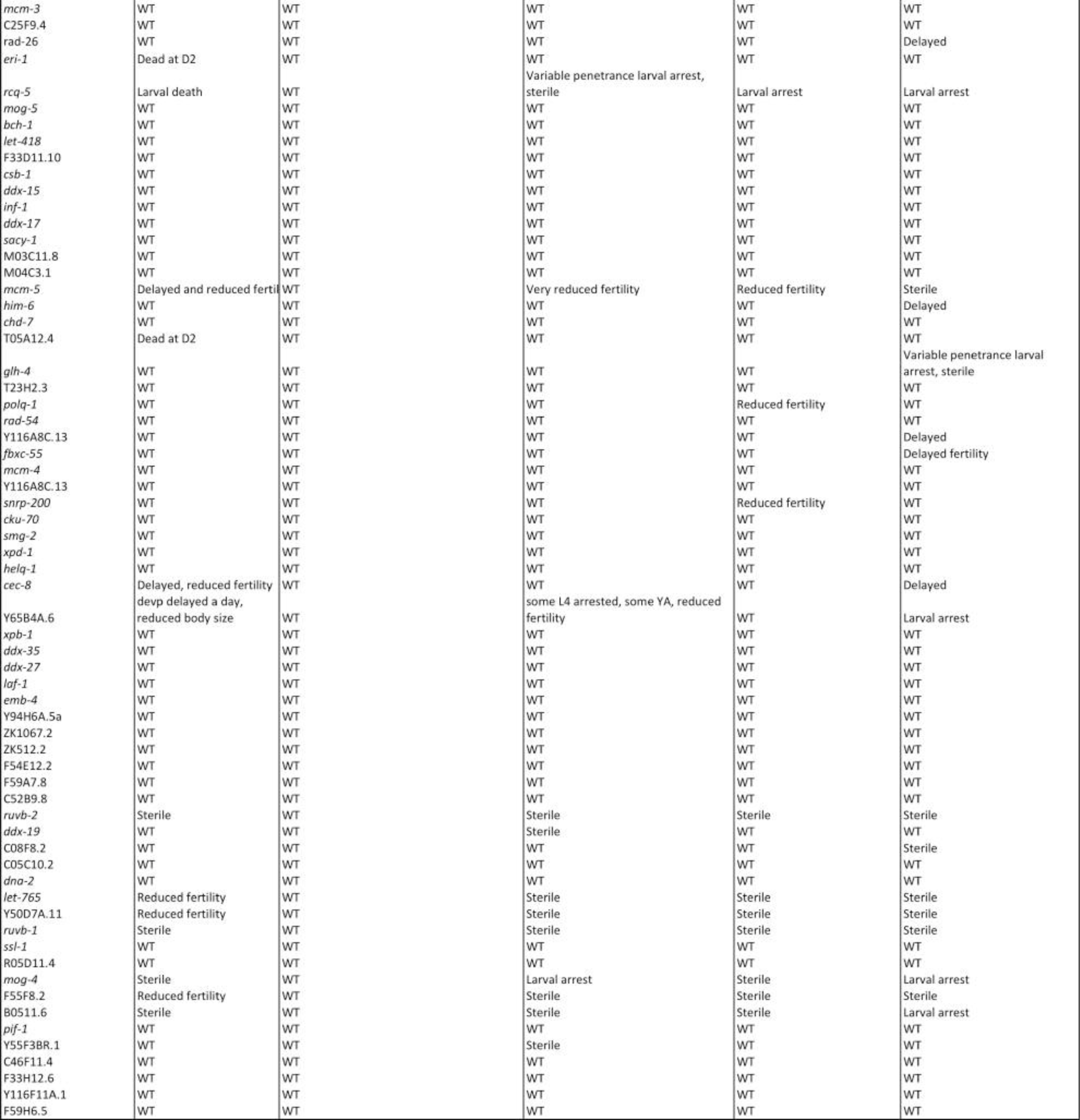
Complete Results of screen for alternate mtDNA replicative helicase. Indicated are gene name or locus, predicted nucleic acid type interacting with the protein, and score (from 0 to 3) for a developmental delay, arrest, and sterility as assessed by microscopy in different genetic backgrounds (iii-vii).

